# Perceptual training selectively strengthens top-down signaling to sensory cortex

**DOI:** 10.64898/2026.06.25.734556

**Authors:** Matheus Macedo-Lima, Mason McCollum, Melissa L. Caras

## Abstract

Sensory cortical neurons adjust their responses to enhance the detection and discrimination of behaviorally relevant stimuli. Top-down signals from frontal cortical regions are thought to mediate these changes, and prior work has proposed that these signals may strengthen during perceptual learning. However, this hypothesis has not been tested directly. Here, we asked whether neurons in the orbitofrontal cortex (OFC), a higher-order brain region implicated in top-down control of sensory cortex, alter their activity during auditory perceptual learning and whether learning-related changes are transmitted to sensory cortex. We found that OFC neurons encode both trial outcome and stimulus-related information, and that the strength and sensitivity of these signals increase as animals learn to detect progressively weaker amplitude modulations. Notably, learning-related changes were conveyed to auditory cortex, but not visual cortex, indicating that perceptual learning selectively strengthens signaling within a task-relevant frontal-to-sensory cortical circuit. Together, these findings provide direct evidence that top-down cortical networks are modified during perceptual learning.

## INTRODUCTION

Practice can refine the ability to perceive sensory stimuli, leading to more sensitive detection thresholds and heightened acuity. This process—termed perceptual learning^1^—occurs across sensory modalities and contributes to complex skills, such as speech comprehension^2–9^ and musical ability^10–16^. However, the neural circuits that enable perceptual learning remain incompletely understood.

Neural recordings consistently demonstrate learning-related changes in the sensitivity or tuning of sensory cortical neurons to trained stimulus features^17–29^. Yet perceptual learning is also strongly influenced by non-sensory factors, including attention, reward, and task demands^30–32^. These factors robustly modulate sensory cortical activity^33–36^, raising the possibility that learning-related plasticity may occur not only within sensory cortex, but also in the top-down brain networks that shape sensory processing.

Indirect support for this idea comes from neurophysiological studies showing that sensory cortical stimulus representations are enhanced when stimuli acquire behavioral relevance, and that this enhancement grows with perceptual learning^19,28,37–39^. These observations are consistent with findings from human neuroimaging workbyer^40,41^ and computational models of perceptual learning^42–44^, and collectively have led to the hypothesis that perceptual learning strengthens top-down signaling to sensory cortex^27^. However, direct evidence for this hypothesis is lacking.

Among the candidate sources of top-down signals to sensory cortex, the orbitofrontal cortex (OFC) is especially compelling because it integrates sensory, value, and task-related information^45–53^, and sends direct projections to multiple sensory cortical areas^54–60^. In the auditory cortex, OFC input modulates sound-evoked responses, conveys task and stimulus-related information, and can drive rapid auditory cortical plasticity^58,59,61^. Consistent with these observations, we recently found that the OFC is required for behavioral sound detection and for task-dependent engagement of auditory cortical neurons^62^. These findings led us to hypothesize auditory perceptual learning enhances OFC-to-auditory cortical communication to enhance the representation of behaviorally relevant sounds.

To test this hypothesis, we monitored OFC neurons and OFC projections to auditory cortex as Mongolian gerbils trained and improved on an auditory detection task. We found that OFC neurons transmit both trial outcome and stimulus-related information to the auditory cortex, and that the strength and sensitivity of these signals increase with learning. Notably, learning-related changes were not observed in OFC projections to visual cortex, indicating that plasticity was restricted to the OFC-to-auditory cortex pathway. Together, these findings provide direct evidence that perceptual learning is accompanied by strengthened signaling within a frontal-to-sensory cortical circuit.

## RESULTS

### Animals learn to detect weak amplitude modulations

To evaluate how orbitofrontal-auditory signaling evolves over the course of perceptual learning, we trained freely-moving Mongolian gerbils to detect amplitude modulations (AM) embedded in broadband noise^63–66^. In this task, water-restricted animals learn to drink steadily from a spout while in the presence of the no-go stimulus (non-AM noise) and to withdraw from the spout when the no-go stimulus smoothly transitions to the go stimulus (5 Hz sinusoidal AM noise). Spout withdrawal is reinforced by pairing the go stimulus with a mild shock (Figure 1A). Animals were trained once per day, every 1-2 days.

**Figure 1.**
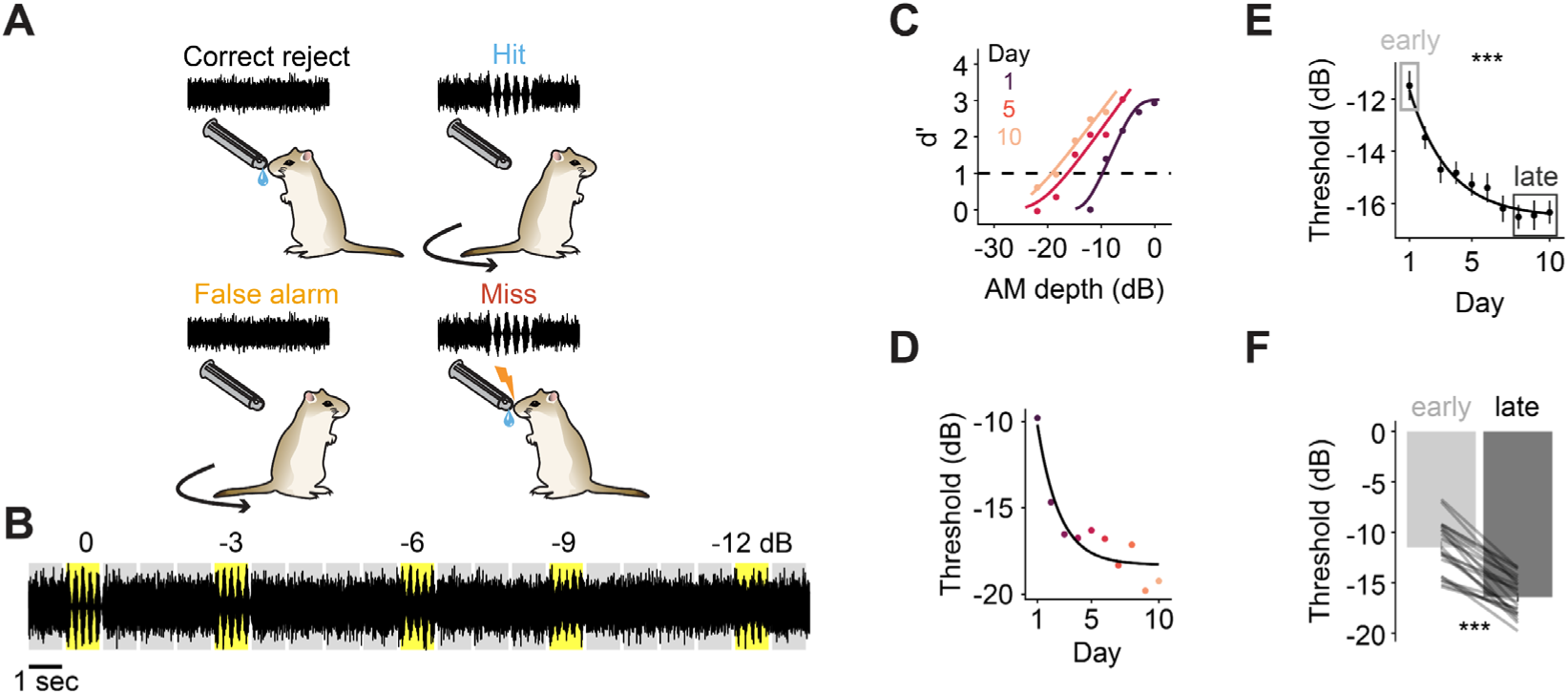
Animals learn to detect weak amplitude modulations. **(A)** Behavioral paradigm. Water-restricted gerbils learned to lick from a waterspout during continuous unmodulated noise and to cease licking when amplitude-modulated (AM) noise was introduced. Failure to withdraw during the AM stimulus resulted in the delivery of a mild shock. **(B)** Representative sound stream from the perceptual training phase. A range of modulation depths bracketing each subject’s current perceptual threshold are presented. Gray shadings indicate non-AM (no-go) trials. Yellow shadings indicate AM (go) trials. **(C)** Psychometric functions from one representative subject on selected perceptual training days (1, 5 and 10). Dashed horizontal line indicates *d’* = 1, used as the detection threshold criterion. **(D)** Detection threshold across training days for the subject in (C), with fitted exponential decay. **(E)** Mean detection thresholds ± SEM across training days for all 22 subjects that successfully learned. Boxes indicate the early and late training stages used throughout the manuscript. **(F)** Thresholds significantly improve from early to late training. Late-stage datapoints represent averages across the final 2-3 perceptual training days. ***p <; 0.001

Animals were initially trained with a single easy go stimulus (0 dB relative to 100% depth AM noise; procedural training) until they reached a predetermined performance criterion (*d’* ≥ 2 on two consecutive training sessions, see STAR Methods). Gerbils achieved this criterion within 49-718 trials (median = 267 trials) across 4-7 sessions (median 4 days) (n = 26 animals across all experiments; 15 females). After reaching criterion, animals transitioned to perceptual training, during which we presented a range of progressively weaker AM depths over the course of ten days (Figure 1B). Figure 1C shows a representative subject’s performance expressed as *d’* as a function of AM depth on three selected days spanning the training period. Threshold was defined as the AM depth at which the psychometric fit crossed *d’* = 1. Figure 1D shows this subject’s threshold across the 10 days of perceptual training, reflecting an improvement rate of 4.92 dB/log_10_(day).

At the group level, thresholds gradually improved across sessions (p < 0.001), at an average rate of 4.90 dB/log_10_(day), and significantly differed between early and late training epochs (Figure 1E-F, p < 0.001). Subjects (n = 4, 3 females) who failed to learn were excluded from analyses involving learning-related changes (see Figure S1A and STAR Methods).

We also measured the slopes and lapse rates of the psychometric functions across training. Psychometric slopes are a proxy for perceptual discrimination precision—i.e., steeper slopes indicate that detection performance transitions sharply from near-zero to near-maximum over a narrow range of AM depths, implying finer sensitivity around threshold. Lapse rates capture the probability of a random failure to respond even to clearly suprathreshold stimuli, independent of perceptual sensitivity. In practice, well-trained animals should show lapse rates near zero, indicating reliable responding at suprathreshold AM depths. In our sample, psychometric slopes significantly increased across training (p = 0.004, Figure S1B), while lapse rates remained low (1.9 ± 0.3%, mean ± SEM) and unchanged (p = 0.725, Figure S1C).

Together, these data confirm that our task induces perceptual learning of AM depth detection.

### OFC neurons encode trial outcome and are sensitive to amplitude modulation depth

We first sought to identify the dominant variables encoded by OFC neurons during performance of our AM detection task. To do so, six animals (four females) underwent procedural training until they reached our performance criterion of *d’* ≥ 2 on two consecutive days. We then implanted these animals with chronic 64-channel silicon probe arrays targeting the left OFC (Figure S2A) and recorded *in vivo* extracellular single-unit activity over the course of perceptual training (Figure 2A). We recorded from a total of 771 single-units and 959 multi-units. Here, we focus on the responses of the 656 single-units classified via k-means clustering as regular-spiking (putative excitatory) neurons (Figure S2B; see STAR Methods), initially pooling data across training days to establish basic response properties.

**Figure 2.**
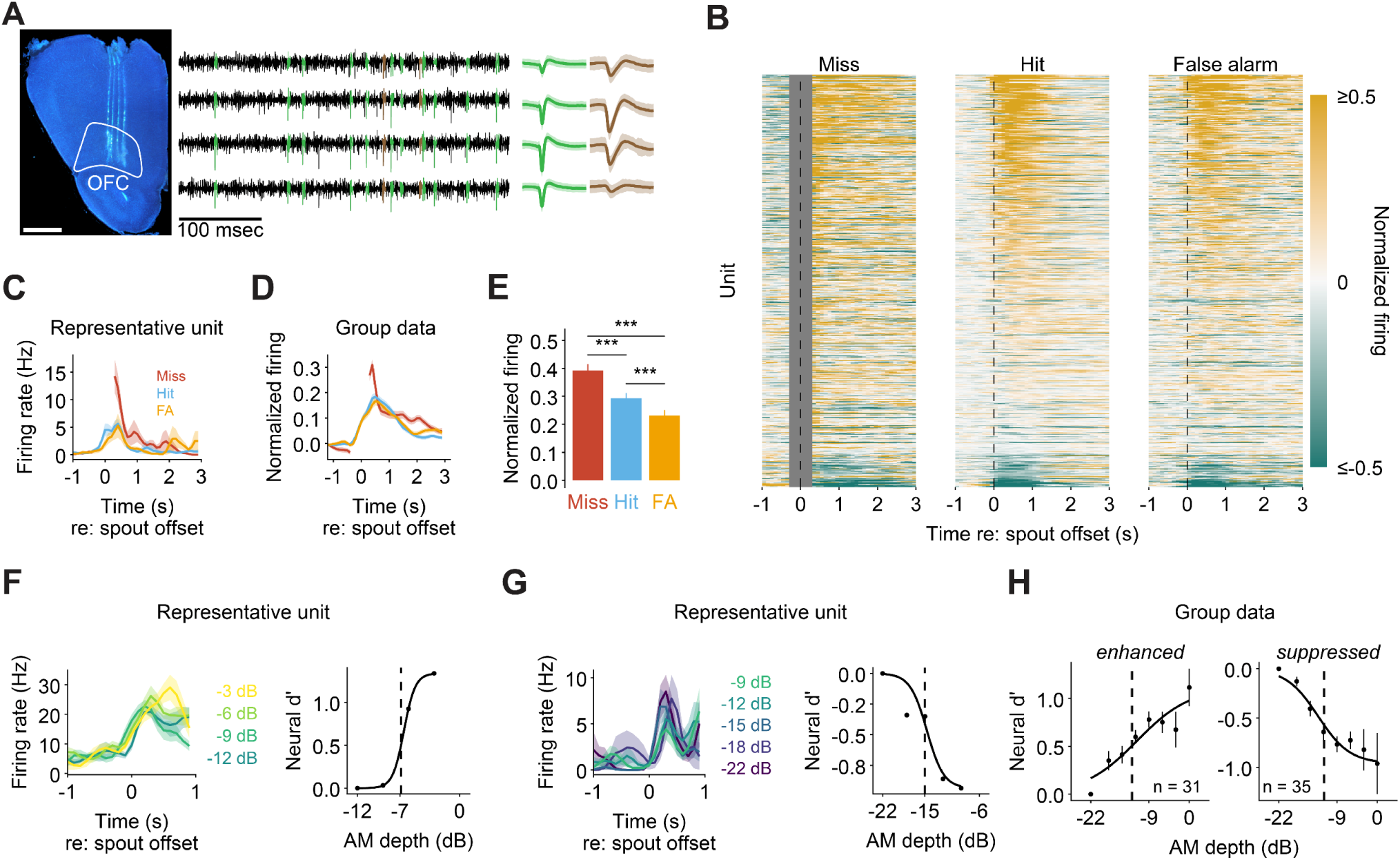
OFC single units encode trial outcome and are sensitive to amplitude modulation depth. **(A)** Extracellular recordings were made using 64-channel silicon arrays targeting the left OFC of freely moving Mongolian gerbils. Electrode placement was confirmed histologically post-hoc via fluorescently labeled tracks and/or electrolytic lesions (left). Scale bar: 1 mm. A representative recording segment (middle) shows high-pass filtered voltage traces from four neighboring channels, with identified single units highlighted in green and brown. Mean ± SD spike waveforms are shown for each isolated unit (right). **(B)** Heatmaps illustrate normalized firing aligned to spout offset (vertical dashed lines). OFC units (rows) are ordered by mean firing between 0 and 1 sec during hit trials, and the same order is used for miss and false alarm trials. Grey region in the miss trial heat map reflects signal blanking due to the shock artifact. **(C)** Mean ± SEM firing of a representative unit during each trial outcome. Firing is aligned to spout offset and the gap in the miss trace (−0.3 to 0.3 sec) reflects signal blanking during shock artifact. **(D)** Mean ± SEM normalized firing across 656 regular-spiking single units, aligned to spout offset. **(E)** Normalized firing quantified within 0-1 sec post-offset (0.3-1.3 sec for misses, avoiding the shock artifact). **(F)** Left: Mean ± SEM firing of a representative unit aligned to spout offset and separated by AM depth. Only hit trial activity is shown. This unit exhibited increased spout offset activity as the AM depth increased. Right: Neural *d’* (relative to the lowest AM depth with ≥ 3 hit trials, -12 dB) as a function of AM depth for the same OFC unit, fit with a sigmoid. The dashed vertical line indicates the neural threshold—the AM depth at which the fit crosses *d’* = 0.5. **(G)** Same as (F) but for a representative unit that exhibited decreased activity as the AM depth increased. Here, threshold is defined as the crossing of *d’* = -0.5. **(H)** Population data from all units exhibiting AM depth sensitivity during the post spout-offset period, separated by the direction of the relationship between neural *d’* and AM depth. Datapoints are means ± SEM. Dashed lines represent the average threshold across individual units. ***p <;0.001.

OFC has been implicated in value encoding, outcome signaling, and reinforcement learning^45,67–70^. We therefore first asked whether log-transformed, baseline subtracted firing (hereafter referred to as “normalized firing”; see STAR methods) differed across trial outcomes in our task (Figure 2B-E). Visualization of firing from an individual unit (Figure 2C) and across the population (Figure 2D) revealed a pronounced increase in activity following spout offset, with elevated activity persisting for up to 3 sec after the event. Spout offsets associated with shock delivery following misses evoked the strongest responses, exceeding those observed on hit and false alarm trials (all p < 0.001; Figure 2E). Responses on hit trials were also greater than those on false alarm trials (p < 0.001).

To determine how widespread outcome-related signaling was across the recorded population, we quantified spout offset responsiveness at the single unit level. Using a permutation test, we compared each unit’s firing during the outcome period to the firing during the immediately preceding non-AM noise epoch. Most units (488/656; 74%) exhibited a significant response to spout offset for at least one trial outcome. Of those, 302/488 (62%) responded during two or three outcome types (Figure S2C-E).

Although prior work has shown that OFC neurons can respond to sounds^46,59,71,72^, including AM cues^58^, most of these studies used high intensity sounds that could drive strong onset responses. We therefore asked whether OFC units are sensitive to the behaviorally-relevant acoustic feature in our task: changes in AM depth embedded within an ongoing, moderate intensity noise carrier. We restricted this analysis to miss trials because during hit trials, spout withdrawals temporally overlap with the AM stimulus, and could therefore confound sensory responses. We found that a small subpopulation of OFC neurons (28/656; 4%) were sensitive to AM depth (see STAR Methods). Consistent with prior reports^59,72^, responses were characterized by slow, sustained activity shifts following sound onset (Figure S2F-G)

OFC neurons can multiplex sensory and behavioral inputs to output value-rich sensory information^46,73,74^. Therefore, we asked whether AM-sensitive units also exhibited spout offset responses. Similar to the overall OFC population (Figure S2C), we found that 68% (19/28) of AM-sensitive units exhibited a significant response to spout offset during at least one trial outcome. Of those, 16/19 (84%) responded during two or three outcome types.

Because the analyses above excluded hit trials—which comprise the majority of strong AM depth presentations—we likely underestimated the prevalence of AM-sensitive OFC neurons. We therefore extended our analysis to ask whether AM depth information was also present during the post-spout offset period on hit trials.

To quantify AM sensitivity during this epoch, and enable comparisons across neurons, we calculated neural *d′* values using responses to the lowest AM depth presented to each unit as the reference condition (see STAR Methods). Conceptually, this metric assesses whether AM depths are distinguishable during the post-offset epoch, with larger *d′* values reflecting greater separation from the weakest AM depth. To identify AM sensitive neurons, we plotted each unit’s *d’* as a function of AM depth and fit the resulting functions with a sigmoid. Units whose fits differed significantly from a flat line and crossed an absolute *d’* of 0.5 were classified as AM depth sensitive (see STAR Methods).

This analysis revealed that approximately 10% of neurons (66/656) exhibited AM depth sensitivity during the post-spout offset period. As illustrated in Figure 2F-H, the relationship between neural *d’* and AM depth could be either positive (31/66) or negative (35/66). AM depth threshold (defined as the depth at which the neurometric fit crossed an absolute *d’* value of 0.5) did not differ between units with positive and negative fits (p = 0.568). Collectively, these findings suggest that a subpopulation of OFC units multiplex AM depth and trial outcome information.

### Perceptual training selectively strengthens outcome signaling in OFC neurons

Our initial analyses revealed that OFC neurons are strongly activated following spout offsets. To determine whether these signals evolve over the course of perceptual training, we used generalized linear models (GLMs) to quantify changes in post-spout offset firing across two training stages: early (day 1) and late (days 8-10). Because a sizeable population of OFC neurons is AM-sensitive, we controlled for AM depth on hit and miss trials by restricting analyses to depths presented in both early and late training for each subject, and by including AM depth as a covariate in the GLM.

As shown in Figure 3, training increased spout offset-driven responses in a trial outcome-dependent manner. Offset-related firing was significantly stronger late in training on hit trials (p = 0.016) but not miss trials (p = 0.601). Responses on false alarm trials also increased late in training (p = 0.008). A day-by-day effect of training on normalized firing was also apparent (Figure S3). A polynomial GLM confirmed that normalized firing rates during hit and false alarm trials increased, particularly in the final days of training (both p = 0.015). In contrast, miss trial firing exhibited a transient decrease that renormalized by late training (p = 0.004). Because both hit and false alarm trials involve voluntary spout withdrawal responses, these findings suggest that perceptual training preferentially enhances OFC signals associated with perceived stimulus detection.

**Figure 3.**
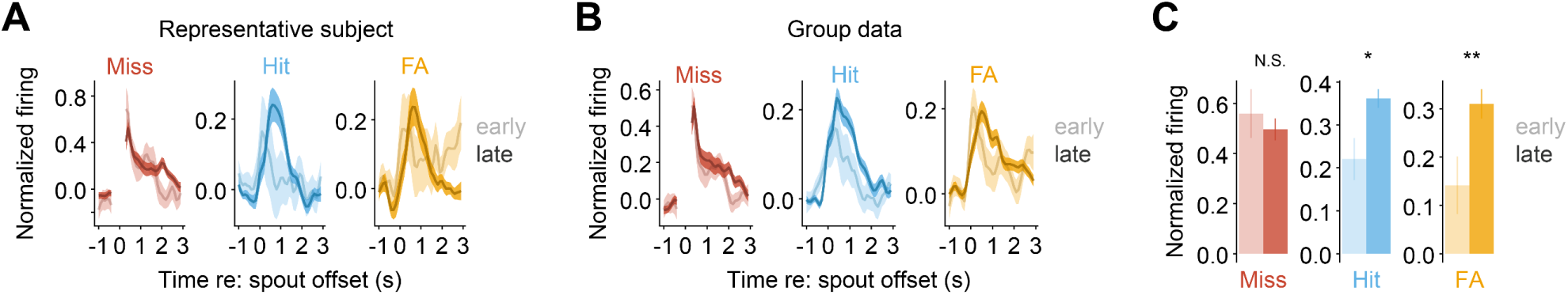
Perceptual training selectively strengthens trial outcome responses in OFC neurons. **(A)** Mean ± SEM normalized firing from a representative subject during each trial outcome during early (n = 15 units) and late training (n = 67 units). Firing is aligned to spout offset and the gap in the miss trace (−0.3 to 0.3 sec) reflects signal blanking during shock artifact. (**B)** Same as A, but for units recorded across all subjects (n = 61 early, 191 late). **(C)** Normalized firing (0-1 sec) increases from early to late training specifically in hit and false alarm trials. *p < 0.05; **p < 0.01.

### Perceptual training improves AM depth sensitivity in OFC neurons

Having established that perceptual training selectively enhances outcome-related signaling in OFC neurons, we next asked whether training also alters AM depth representations during the post-spout offset period. To address this question, we focused on the 10% (66/656) of OFC neurons that exhibited AM sensitivity during this epoch on hit trials.

Figure 4A depicts spout-offset related activity separated by AM depth for two representative neurons recorded during early (top) and late training (bottom). The neuron recorded during early training exhibited graded but modest responses to strong AM depths, resulting in a relatively poor neural threshold (−5 dB, top right). In contrast, the neuron recorded during late training shows stronger responses at lower depths, resulting in a substantially lower threshold (−13 dB, bottom right). This pattern was recapitulated at the population level (Figure 4B-C), such that on average, neural thresholds were significantly lower (better) late in training (GLM/ANOVA; p < 0.001; Figure 4D).

**Figure 4.**
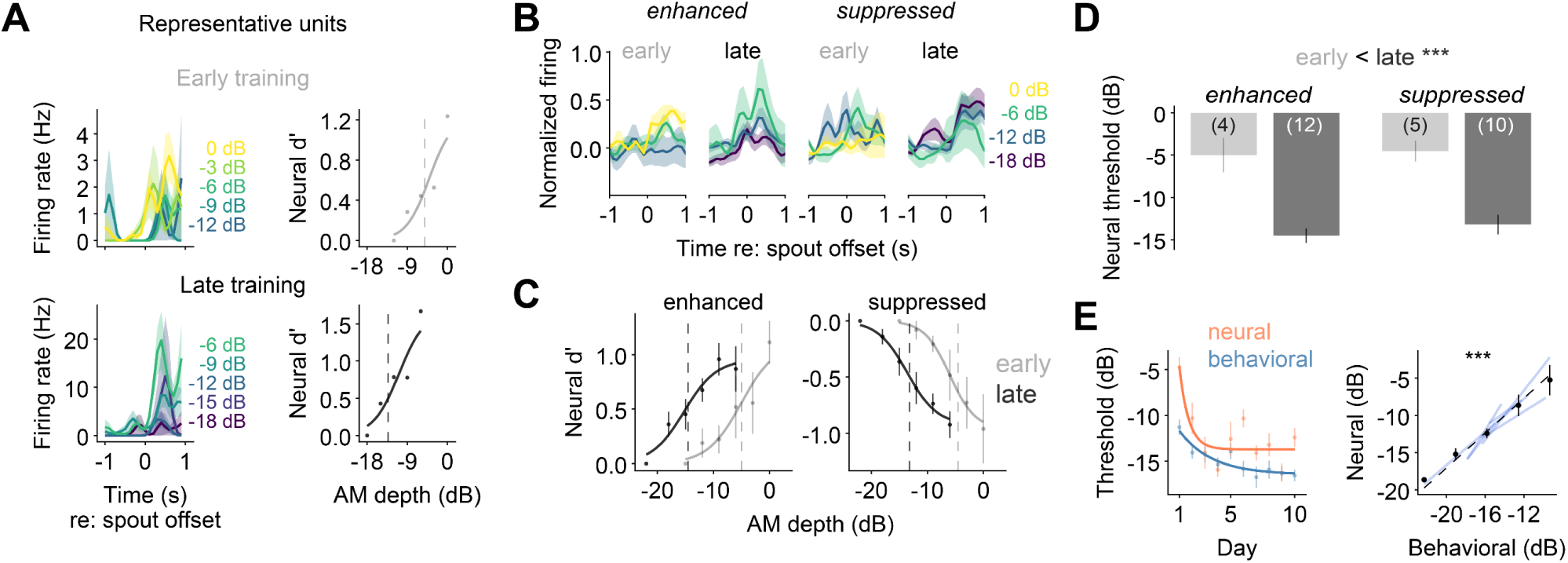
Perceptual training improves AM depth sensitivity in OFC neurons. **(A)** Representative examples of two units from early (top) and late training (bottom). Mean ± SEM firing rates aligned to spout offset on hit trials (left) show clear AM depth sensitivity in both examples, but an increased response to weaker AM depths in the late training example (bottom). Neural *d’* and neurometric fits (right) reveal a lower (better) threshold (dashed line) in the late training example (bottom). **(B)** Mean ± SEM normalized firing rates across the population of neurons exhibiting AM depth sensitivity during the post spout-offset period, separated by training stage and by the direction of the relationship between neural *d’* and AM depth. **(C)** Neurometric fits during early and late training for the population of neurons included in panel (B). Datapoints are means ± SEM. Dashed lines represent the average threshold across individual units for each training stage. **(D)** Neural thresholds significantly improve from early to late training. Sample sizes are indicated within bar plots. **(E)** Left: Both behavioral and neural thresholds significantly improve across training. Datapoints are means ± SEM across subjects (n = 6) or units (n = 66), respectively. Lines indicate exponential fits. Right: Neural and behavioral thresholds are strongly correlated. Datapoints are means ± SEM across units (n = 66) and subjects (n = 6), binned into 5 behavioral threshold ranges. Dashed line indicates overall linear fit across all data. Blue solid lines depict fits for individual subjects. ***p <;0.001.

To determine whether neural and behavioral sensitivity improve in parallel over training, we plotted thresholds as a function of training day (Figure 4E, left). Both neural and behavioral thresholds significantly decreased across training (p < 0.001, GLM), indicating improved sensitivity. Improvements occurred in tandem, resulting in a strong positive relationship between neural and behavioral thresholds (p < 0.001, GLM). Figure 4E (right) illustrates the overall relationship along with individual subject fits (blue lines).

Collectively, these findings demonstrate that OFC neurons multiplex AM depth and trial outcome-related information, and that these signals grow stronger over perceptual learning.

### Perceptual training selectively strengthens outcome signaling in the OFC-to-AC pathway

OFC neurons are functionally heterogenous^46^ and project to numerous downstream targets^55,75^. To determine whether trial outcome and AM depth information are represented in OFC projections to auditory cortex (AC) during our task, we virally expressed the genetically encoded calcium indicator jGCaMP8s in OFC axons of nine animals (four females; Figure 5A; Figure S4A). We then used fiber photometry to record calcium signals from OFC axons in left AC across 10 days of perceptual training (Figure 5B).

**Figure 5.**
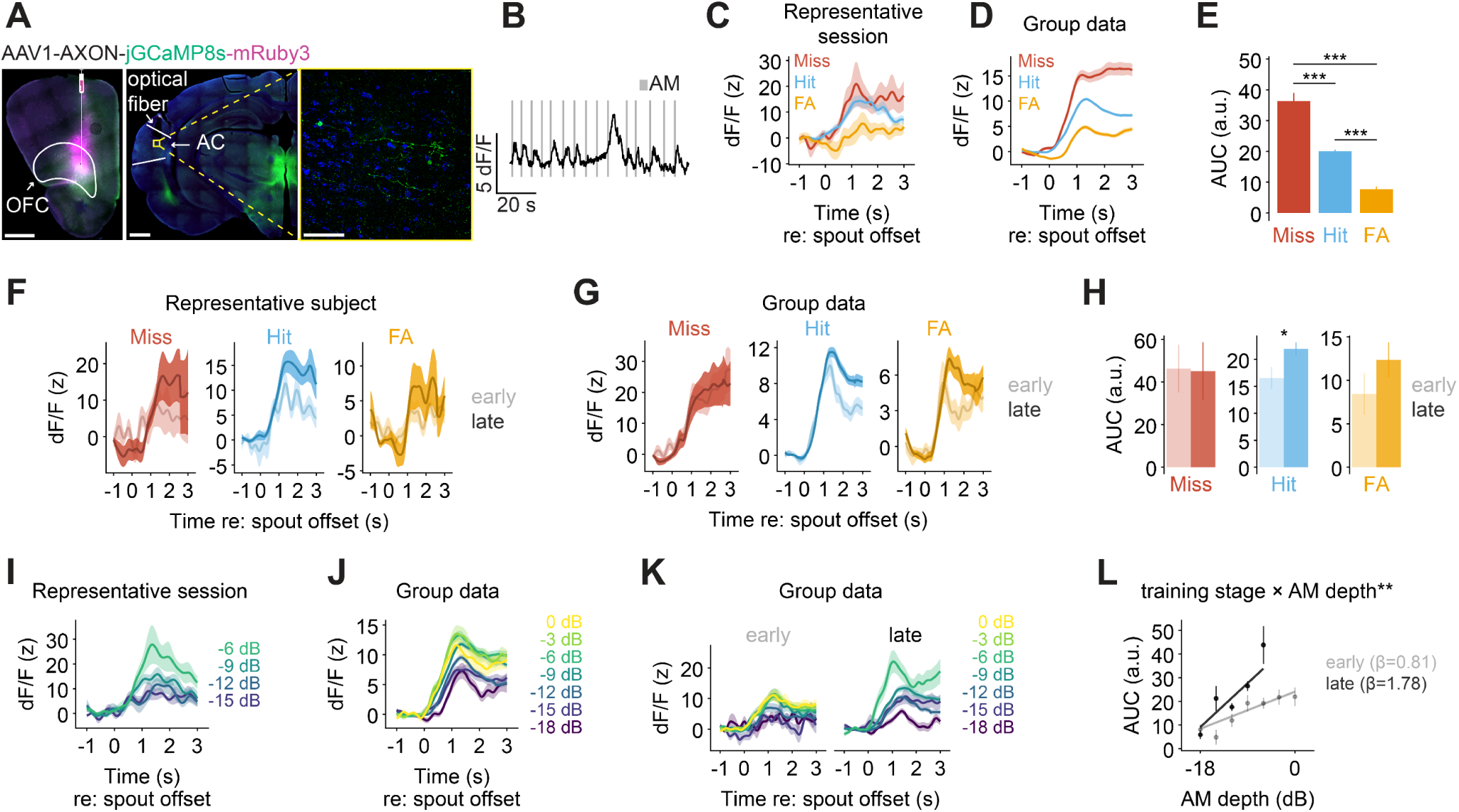
Perceptual training selectively strengthens outcome signaling and improves AM depth sensitivity in the OFC-to-auditory cortex pathway. (A) AAV1-AXON-jGCaMP8s-mRuby3 was injected into OFC and an optical fiber was implanted in auditory cortex (AC) to record GCaMP signals from OFC axons. Left: coronal section showing injection site in OFC. Middle: coronal section showing fiber placement above AC with axonal labeling (green). Right: magnified view of the fiber tip region (yellow dashed box) showing GCaMP8s-expressing OFC axons within AC. Scale bars: 1 mm (left and middle) and 50 µm (right). (B) Representative trace from one animal during task performance. Gray ticks indicate AM stimulus delivery. (C) Representative single-session signal (mean ± SEM) aligned to spout offset during each trial outcome type. (D) Same as (C) but averaged across all animals (n = 9) and sessions (n = 84). (E) The area under the curve (AUC; 0-3 sec post-spout offset) was quantified for each trial outcome type. Data are means ± SEM across all animals and sessions. (F) Mean ± SEM calcium signals from a representative subject during early and late training, aligned to spout offset for each trial outcome type. (G) Same as (F) but pooled across all animals. (H) Mean ± SEM AUC values separated by trial outcome type and training stage. (I) Representative single-session signal (mean ± SEM) aligned to spout offset during hit trials and separated by AM depth. (J) Same as (I) but averaged across all subjects and sessions. (K) Population-averaged signals (mean ± SEM) aligned to spout offset during hit trials separated by AM depth and by training stage. (L) AUC as a function of AM depth during early (gray) and late (black) training stages. Slopes of the linear fits (AUC units/dB) are indicated in the panel legend. **p <; 0.01; ***p <; 0.001.

Visualization of normalized calcium signals pooled across training days revealed robust responses following spout offset, with elevated activity persisting for several seconds (Figure 5C-D). To quantify the magnitude of these signals, we calculated the area under the curve (AUC) during the three seconds following spout offset. Consistent with our electrophysiological results, spout offsets associated with shock delivery following misses evoked the largest AUC values, followed by hits, then false alarms (Figure 5E; all p < 0.001).

Perceptual learning is associated with enhanced outcome signaling in OFC putative projection neurons (Figure 3). To determine whether OFC conveys these learning-related changes to AC, we compared calcium signals from OFC axons across early and late training. For this and all subsequent training-related analyses of OFC-auditory signaling, we excluded one subject that failed to exhibit perceptual learning (see STAR Methods and Figure S1A), yielding a final sample of eight subjects (four females).

Figure 5F shows spout offset-aligned calcium signals during early and late training for one representative subject. In this example, training-related changes are evident selectively on hit trials. Group level analyses that controlled for AM depth revealed that training increased spout-offset related signals on hit trials (p = 0.028), with no effect on miss trials (p = 0.536) or false alarm trials (p = 0.934; Figure 5G-H). We also used GLMs to test the day-by-day effect of training on AUC values. As expected, AUC values on hit trials increased during training (p = 0.024), whereas those on false alarm trials (p = 0.944) and miss trials (p = 0.113) remained unchanged (Figure S4B). Together, these findings indicate that perceptual learning selectively strengthens outcome-related signaling in the OFC-to-AC pathway.

### Perceptual training improves AM depth sensitivity in the OFC-to-AC pathway

Because OFC neurons multiplex AM depth and outcome information during the post-spout offset period, we next asked whether AM depth information was similarly present in the OFC projections to AC. To address this question, we aligned calcium signals to spout offset on hit trials and separated trials by AM depth. On average, signals scaled with modulation strength (Figure 5I-J). To quantify this effect, we used a GLM to test whether AM depth predicted the area under the curve (AUC; 0-3 sec post-spout offset), while controlling for training day. AM depth strongly predicted AUC magnitude (p < 0.001; Figure S4C). Similar AM-dependent scaling was observed at the level of individual training sessions, with 51% of sessions (43/84) exhibiting significant positive correlations between AM depth and AUC (Figure S4D).

Perceptual training enhances AM depth sensitivity during the trial outcome period in a subset of OFC neurons (Figure 4). To determine whether similar changes are reflected in OFC projections to the auditory cortex, we compared spout offset-aligned calcium signals across training stages. On the first day of training, calcium signals largely overlapped across AM depth, indicating weak sensitivity (Figure 5K; left). By late training, signals exhibited a clearer separation across AM depths (Figure 5K; right).

To quantify this effect, we modeled AUC values (0-3 sec post-spout offset) as a function of AM depth and training stage. AUC significantly increased with AM depth during both early (p = 0.001) and late (p < 0.001) training, but the relationship was significantly steeper late in training (interaction p = 0.008; Figure 5L). These results suggest that perceptual training strengthens AM depth sensitivity in the OFC-to-AC pathway.

### Auditory perceptual training does not alter signaling in the OFC-to-visual cortex pathway

To determine whether training-related changes are specific to OFC-auditory signaling or reflect a global change in OFC output, we examined the OFC projection to primary visual cortex (VC), a region thought to be minimally involved in auditory detection^76–78^. We virally expressed the genetically encoded calcium indicator jGCaMP8s in OFC axons of 11 animals (seven females; Figure 6A, Figure S5A) and used fiber photometry to record calcium signals from OFC axons in left VC across 10 days of auditory perceptual training (Figure 6B).

**Figure 6.**
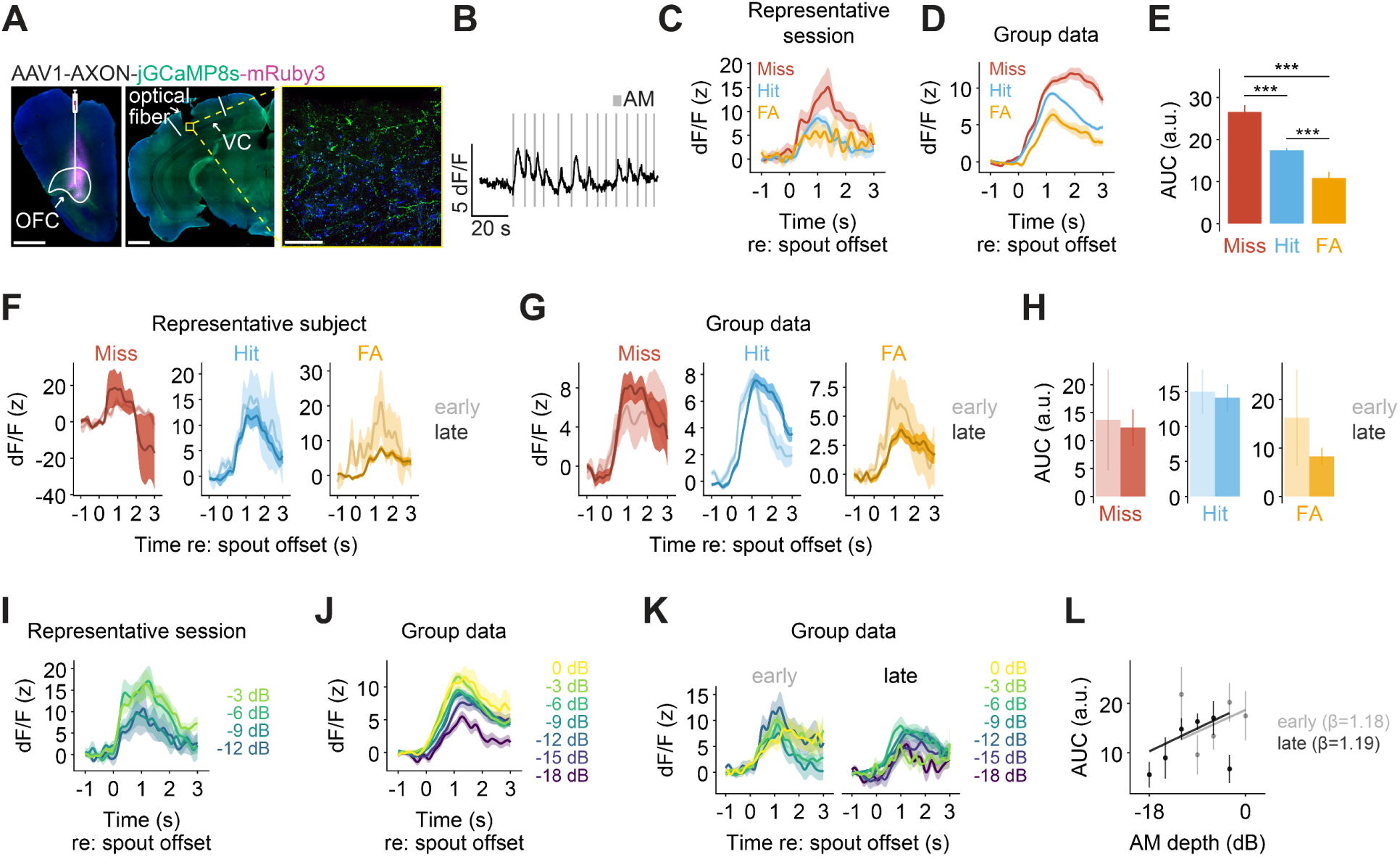
Auditory perceptual training does not alter signaling in the OFC-to-visual cortex pathway. **(A)** AAV1-AXON-jGCaMP8s-mRuby3 was injected into OFC and an optical fiber was implanted in visual cortex (VC) to record GCaMP signals from OFC axons. Left: coronal section showing injection site in OFC. Middle: coronal section showing fiber placement above VC with axonal labeling (green). Right: magnified view of the fiber tip region (yellow dashed box) showing GCaMP8s-expressing OFC axons within VC. Scale bars: 1 mm (left and middle) and 50 µm (right). **(B)** Representative trace from one animal during task performance. Gray ticks indicate AM stimulus delivery. **(C)** Representative single-session signal (mean ± SEM) aligned to spout offset during each trial outcome type. **(D)** Same as (C) but averaged across all animals (n = 11) and sessions (n = 108). **(E)** The area under the curve (AUC; 0-3 sec post-spout offset) was quantified for each trial outcome type. Data are means ± SEM across all animals and sessions. **(F)** Mean ± SEM calcium signals from a representative subject during early and late training, aligned to spout offset for each trial outcome type. **(G)** Same as (F) but pooled across all animals. **(H)** Mean ± SEM AUC values separated by trial outcome type and training stage. **(I)** Representative single-session signal (mean ± SEM) aligned to spout offset during hit trials separated by AM depth. **(J)** Same as (I) but averaged across all subjects and sessions. **(K)** Population-averaged signals (mean ± SEM) aligned to spout offset during hit trials separated by AM depth and by training stage. **(L)** AUC as a function of AM depth during early (gray) and late (black) training stages. Slopes of the linear fits (AUC units/dB) are indicated in the panel legend. ***p <; 0.001.

**Table 1.** Statistical results.

| Figure | Test | Predictor(s) | Statistic | P value |
| --- | --- | --- | --- | --- |
| 1E | GLM/ANOVA | Log<sub>10</sub>(Day) | $\chi^2_1 = 62.520$ | **< 0.001** |
| 1F | GLM/ANOVA | Training stage | $\chi^2_1 = 132.260$ | **< 0.001** |
| 2E | GLM/ANOVA | Trial outcome | $\chi^2_2 = 95.152$ | **< 0.001** |
|  | Bonferroni: Trial outcome | Miss – Hit | Z = 5.815 | **< 0.001** |
|  |  | Miss – FA | Z = 9.678 | **< 0.001** |
|  |  | Hit – FA | Z = 3.971 | **< 0.001** |
| 3C | GLM/ANOVA: Miss | Training stage | $\chi^2_1 = 0.274$ | 0.601 |
| | | AM depth (covariate) | $\chi^2_1 = 1.891$ | 0.169 |
| | GLM/ANOVA: Hit | Training stage | $\chi^2_1 = 5.756$ | **0.016** |
| | | AM depth (covariate) | $\chi^2_1 = 1.029$ | 0.310 |
| | GLM/ANOVA: FA | Training stage | $\chi^2_1 = 6.985$ | **0.008** |
| 4D | GLM/ANOVA | Training stage | $\chi^2_1 = 53.995$ | **< 0.001** |
| | | Modulation direction | $\chi^2_1 = 0.530$ | 0.467 |
| | | Interaction | $\chi^2_1 = 0.117$ | 0.733 |
| 4E | GLM/ANOVA | Behavioral threshold | $\chi^2_1 = 51.885$ | **< 0.001** |
| 5E | GLM/ANOVA | Trial outcome | $\chi^2_2 = 640.813$ | **< 0.001** |
|  | Bonferroni: Trial outcome | Miss – Hit | Z = 9.377 | **< 0.001** |
|  |  | Miss – FA | Z = 22.365 | **< 0.001** |
|  |  | Hit – FA | Z = 20.583 | **< 0.001** |
| 5H | GLM/ANOVA: Miss | Training stage | $\chi^2_1 = 0.382$ | 0.536 |
| | | AM depth (covariate) | $\chi^2_1 = 0.474$ | 0.491 |
| | GLM/ANOVA: Hit | Training stage | $\chi^2_1 = 4.800$ | **0.028** |
| | | AM depth (covariate) | $\chi^2_1 = 49.568$ | **< 0.001** |
| | GLM/ANOVA: FA | Training stage | $\chi^2_1 = 0.007$ | 0.934 |
| 5L | GLM/ANOVA | Training stage | $\chi^2_1 = 15.571$ | **< 0.001** |
| | | AM depth | $\chi^2_1 = 58.472$ | **< 0.001** |
| | | Interaction | $\chi^2_1 = 7.085$ | **0.008** |
| 6E | GLM/ANOVA | Trial outcome | $\chi^2_2 = 132.160$ | **< 0.001** |
|  | Bonferroni: Trial outcome | Miss – Hit | Z = 7.965 | **< 0.001** |
|  |  | Miss – FA | Z = 11.471 | **< 0.001** |
|  |  | Hit – FA | Z = 6.458 | **< 0.001** |
| 6H | GLM/ANOVA: Miss | Training stage | $\chi^2_1 = 1.649$ | 0.199 |
| | | AM depth (covariate) | $\chi^2_1 = 2.746$ | 0.097 |
| | GLM/ANOVA: Hit | Training stage | $\chi^2_1 = 0.137$ | 0.711 |
| | | AM depth (covariate) | $\chi^2_1 = 2.500$ | 0.114 |
| | GLM/ANOVA: FA | Training stage | $\chi^2_1 = 0.000$ | 0.998 |
| 6L | GLM/ANOVA | Training stage | $\chi^2_1 = 1.528$ | 0.216 |
| | | AM depth | $\chi^2_1 = 11.836$ | **< 0.001** |
| | | Interaction | $\chi^2_1 = 0.489$ | 0.484 |

GLM: Generalized linear model; ANOVA: Analysis of Variance; FA: False alarm; Bolded p-values indicate statistical significance.

To characterize the signals carried by this projection, we initially pooled calcium signals across animals and training days. Spout offset elicited robust responses (Figure 6C-D), with dynamics similar to those observed in the OFC-to-AC projection: activity peaked after ∼1 sec after spout offset and remained elevated for several seconds. Trial outcome discrimination was also evident, such that spout offsets associated with shock delivery following misses evoked the strongest responses, followed by hits, then false alarms (Figure 6E; all p < 0.001).

To assess whether these signals evolve with experience, we compared calcium signals from OFC axons in VC across early and late training. For these analyses, we excluded three subjects that failed to exhibit perceptual learning (see STAR Methods and Figure S1A), yielding a final sample of eight subjects (four females). Figure 6F depicts spout-offset aligned calcium signals for one representative subject, separated by trial outcome and training stage. Responses largely overlapped across training stages. Group level analyses that controlled for AM depth revealed that response magnitude (AUC) did not differ between early and late training for any trial outcome (all p > 0.232; Figure 6G-H). We also used GLMs to test the day-by-day effect of training on AUC values. Training day had no effect on AUC magnitude for any trial outcome (all p > 0.088; Figure S5B). These findings suggest that auditory perceptual training does not substantially modify outcome-related signaling in the OFC-to-VC pathway.

We next asked whether AM depth information is present in the OFC-to-VC projection during the trial outcome period on hit trials. Similar to OFC-to-AC axons, visual cortical projection activity scaled with AM depth, increasing with modulation strength (Figure 6I-J). To quantify this effect, we used a GLM to test whether AM depth predicted the AUC 0-3 sec post-spout offset, while controlling for training day. AM depth predicted AUC magnitude (p < 0.001; Figure S5C), but AM-dependent scaling was observed in only 23% (25/108; 24/25 positive slopes) of individual training sessions (Figure S5D), a substantially lower fraction than that observed in OFC-to-AC recordings (51%; Figure S4D). A GLM comparing the AM depth × AUC relationship between the OFC-to-AC and OFC-to-VC datasets revealed a significantly stronger relationship (i.e. a steeper AM depth x AUC slope) in OFC-to-AC axons (p < 0.001; Figure S5E). These results suggest that while AM depth information is present in OFC axons within visual cortex, AM depth representations are weaker and less consistently expressed than those observed in auditory cortex.

Finally, to test whether AM sensitivity in the OFC-to-VC pathway is affected by training, we compared OFC axonal responses in the visual cortex across training stages. Calcium signals moderately scaled with AM depth during both early and late training (Figure 6K), and the relationship between AUC magnitude and AM depth did not change from early to late training stages (Figure 6L; p = 0.484).

Together, these findings demonstrate that the OFC-to-VC projection encodes both trial outcome and AM depth, but these signals are not modified by auditory perceptual training.

## DISCUSSION

Decades of work have established that perceptual learning is accompanied by changes in sensory cortical representations of trained stimulus features^19,22–29,37–41,79^. Top-down signals from frontal cortical regions are known modulators of sensory cortical activity^54,56,57,59,61,62,80,81^, and theoretical^42–44^ and experimental work^19,27,28,37–41,79,82–84^ have led to the hypothesis that these signals strengthen with learning. Here we provide direct evidence for this hypothesis by demonstrating that perceptual learning enhances outcome- and stimulus-related signaling in the OFC, and that these learning-related changes are conveyed to OFC axons in auditory cortex, but not visual cortex. Together, our findings suggest that auditory perceptual learning selectively strengthens OFC-to-auditory cortical communication to enhance the representation of behaviorally relevant sounds.

### Functional implications of enhanced OFC-to-AC signaling

We found that most OFC neurons were strongly activated following spout offset. Response magnitudes varied systematically across trial outcomes, indicating that OFC neurons encode task-related information beyond spout withdrawal alone. The strongest responses were observed during miss trials, consistent with OFC’s prominent role in supporting reinforcement-related^50,85^ and prediction error signaling^70^. Notably, responses were also larger on hit trials than on false alarm trials. This difference may reflect variation in the strength of the underlying sensory evidence and/or perceptual confidence^86–88^. Together, these findings are in agreement with decades of work establishing the OFC as a central hub for value and outcome processing^45–53^, and add to a growing body of literature implicating the OFC in outcome computation during sound-guided behavior^58,62,86,89,90^.

Current models of OFC function propose that outcome signals can serve as top-down ‘teaching signals’ that enhance sensory cortical representations of behaviorally-relevant stimuli^73^. Consistent with this framework, we found that outcome-related signals are transmitted from OFC to AC, a region known to exhibit perceptual learning-related improvements in AM sensitivity^19^, and that perceptual learning selectively strengthened hit-related signaling within this pathway. These results suggest that perceptual learning progressively strengthens OFC representations associated with stimulus detection, and that the transmission of these signals to the AC may reinforce or stabilize behaviorally relevant sensory representations, ultimately improving the detection of weak AM cues. Future loss-of-function experiments targeting the OFC-to-AC pathway will be required to directly test this interpretation.

In addition to exhibiting robust spout offset responses, we found that a subpopulation of OFC neurons multiplex information related to AM depth. Notably, these signals exhibit weak AM depth sensitivity early on that strengthens as training progresses, in parallel with behavior. While these changes may simply reflect training-related refinements in ascending acoustic input, we cannot rule out the possibility that they reflect changes in perceived stimulus salience^91^ or perceptual confidence^86–88^. Regardless, our fiber photometry experiments revealed that these signals are fed back to the AC, where they may serve to further boost or refine responses to the behaviorally-relevant AM stimulus ^59,61,73^.

At first glance, our findings may appear inconsistent with recent work showing that OFC activity increases during passive auditory habituation and that OFC silencing reverses habituation-related changes in AC^80^. However, our study fundamentally differs in behavioral context. Whereas habituation reduces the behavioral relevance of repeated sounds, perceptual learning increases the detectability of behaviorally relevant sensory features. Taken together, both studies suggest that OFC adaptively reshapes auditory cortical representations according to behavioral demands, suppressing responses to irrelevant stimuli while enhancing responses to relevant ones. Thus, OFC may act to dynamically sculpt sensory representations based on their relevance for behavior.

### Target specificity of training-related changes in OFC signaling

Although OFC conveys both outcome-related and AM-depth signals to the visual cortex during auditory task performance, neither signal changed with perceptual learning. This result strengthens the argument that OFC may broadcast task state information across the brain to guide behavior^74^, while selectively strengthening information transfer to the sensory cortical area directly involved in solving the task at hand^54,58,62^.

It remains unknown whether improvements in AM depth detection generalize to analogous perceptual judgements in other sensory modalities (such as detecting weak modulations of light intensity). However, some forms of perceptual learning, particularly temporal interval discrimination, can transfer across sensory modalities^92–96^. An intriguing open question is therefore whether auditory temporal interval discrimination learning induces similar plasticity in OFC projections to both auditory and visual cortex. If so, such a finding would suggest that differential recruitment of OFC-to-sensory cortical pathways contributes to the variability in perceptual learning transfer observed across tasks, and may ultimately inform strategies for promoting learning generalization.

## RESOURCE AVAILABILITY

### Lead contact

Further information and requests for resources and reagents should be directed to and will be fulfilled by the Lead Contact, Melissa L. Caras.

### Materials Availability

This study did not generate new unique reagents.

### Data and code availability

All relevant MATLAB, Python and R code for processing behavioral and physiology data are available at https://github.com/caraslab.

## ACKNOWLEDGMENTS

We thank members of the Caras Lab for helpful feedback and assistance with animal care. Special thanks to Dr. Daniel Stolzberg for software assistance. We acknowledge the Imaging Core Facility in the department of Cell Biology and Molecular Genetics at the University of Maryland, College Park for training and the use of the Zeiss LSM 980 Airyscan 2 confocal microscope. Purchase of the Zeiss LSM 980 Airyscan 2 was supported by Award Number 1S10OD025223-01A1 from the National Institute of Health (NIH). This work was supported by NIH R00DC016046 and R01DC020742 to MLC.

## AUTHOR CONTRIBUTIONS

MML and MLC designed research; MML and MM performed research; MML analyzed data; MML and MLC wrote the paper.

## DECLARATION OF INTERESTS

The authors declare no competing interests.

## DECLARATION OF GENERATIVE AI AND AI-ASSISTED TECHNOLOGIES IN THE WRITING PROCESS

During the preparation of this work, the authors used Claude (Haiku, Sonnet, and Opus; Anthropic) and ChatGPT-5.5 (OpenAI) for assistance in data analysis and visualization code and manuscript proofreading. After using these tools, the authors reviewed and edited the content as needed and take full responsibility for the content of the published article.

## STAR METHODS

### EXPERIMENTAL MODEL AND SUBJECT DETAILS

#### Subjects

Adult Mongolian gerbils (*Meriones unguiculatus*; n = 26, 16 females) used in this study were raised from commercially obtained breeder pairs (Charles River). Animals were housed with one or more same-sex littermates under a 12/12 h light/dark cycle and provided with *ad libitum* food and enrichment. Water access was *ad libitum* until animals were placed on controlled water access for behavioral experiments. All procedures were approved by the Institutional Animal Care and Use Committee at the University of Maryland College Park.

## METHOD DETAILS

### Behavioral task and set up

Animals were trained on an aversive go/no*-*go AM detection task as previously described^19,20,63,97^. A custom-designed test arena (CCMI Plastics) containing a stainless-steel water spout and stainless-steel floor plate was placed in the center of a sound attenuating booth (GretchKen). Infrared beam-break detection of the animal snout near the spout triggered water delivery via a syringe pump (NE-1000, New Era Pump Systems). Stimuli were delivered via a calibrated free-field speaker (DX25TG59-04 1” Fabric Dome Tweeter, Peerless by Tymphany) positioned directly above the test cage. Electrical shocks were delivered via an H13-15 Precision Animal Shocker (Colbourn) gated by an H13-XX-SP01 electrical grounding relay (Colbourn) to minimize electrical noise during electrophysiological recordings. Behavior was monitored remotely via a Logitech C270 USB webcam. Data acquisition, sound delivery, and digital outputs were controlled via an RZ6 multifunction processor (Tucker Davis Technologies, TDT) and the ePsych MATLAB toolbox (https://github.com/dstolz/epsych).

### Behavioral training and analysis

Animals first learned to drink steadily from a spout while in the presence of a continuous unmodulated broadband noise (0.1-20 kHz, 45 dB SPL). Animals then underwent associative training, during which they learned to withdraw from the spout when the sound changed to AM noise (5 Hz rate, 100% modulation depth, 1 second duration) to avoid a mildly aversive shock (0.5-1 mA, 300 msec).

No-go trials were defined as discrete 1-sec windows during the continuous non-AM noise stream. Go trials consisted of 1-sec presentations of AM noise. Go trials were pseudorandomly interspersed with 3-5 no-go trials. Trials were only triggered when animals made contact with the water spout.

The animal’s spout contact was evaluated during the last 100 msec of each trial. If the animal broke spout contact for >50 msec during this monitoring window, the response was scored as a spout withdrawal. Spout withdrawal was considered a ‘hit’ on go trials and a ‘false alarm’ on no-go trials. Maintaining spout contact was considered a ‘miss’ on go trials and a ‘correct reject’ on no-go trials (Figure 1A). Miss trials were punished with a shock. Shock intensity was initially set low and adjusted individually due to variability in pain sensitivity across subjects^98^.

Behavioral performance was quantified using the signal detection metric *d’*^99^:

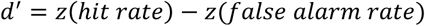

Hit and false alarm rates were set to a floor of 5% and a ceiling of 95% to avoid infinite *d’* values.

Animals initially learned the association by training with a single, fully modulated (100% depth) AM stimulus. Following task acquisition (*d’* ≥ 2 on two consecutive days), animals transitioned to *perceptual training*^19^. During perceptual training, the task was made progressively more challenging by reducing the AM depth across sessions. Because the decision axis for AM detection is logarithmic^100^, depths are typically referred to on a dB scale relative to 100% depth. We adopt that convention here. Thus, 0 dB refers to fully modulated (100% depth) noise, and negative numbers refer to shallower depths. This dB scale should not be confused with dB SPL, which indicates the root-mean-squared (RMS) sound level of the stimulus. To avoid confounding loudness perception with depth perception, the gain of the AM signal was adjusted to control for changes in average power across AM depths^101^.

Each perceptual training session began with the presentation of 0 dB ‘reminder’ trials to ensure the animal was willing to perform the task. Once the animal achieved three consecutive hits or consumed 0.5 mL of water (whichever came first), we began delivering no-go and go trials as described above.

During the first perceptual training session, animals were initially presented with five AM depths selected to bracket the expected detection threshold based on prior literature (0 to -12 dB re: 100% AM, in -3 dB steps)^19,20,63,97,102^. The starting depths in subsequent sessions depended on the animal’s threshold estimate from the previous session. If needed, stimulus values were increased or decreased once per session using predetermined step sizes to maintain threshold bracketing. Perceptual training sessions lasted until animals were satiated or until ∼100 AM trials were completed, whichever happened first. Across all sessions, animals completed a median of 100 AM trials/session (range: 33-109).

To maintain threshold bracketing throughout perceptual training, some AM depths below the animal’s estimated detection threshold were presented. Punishing animals for failing to detect these sub-threshold stimuli would likely increase false alarm rates and/or reduce task engagement. Therefore, sub-threshold AM depths were never paired with a shock. Previous studies have shown that this approach does not prevent perceptual learning^19,20,63,97^ and does not result in secondary conditioning to the presence or absence of shock^103^. For response-aligned neural analyses, only hits, false alarms and suprathreshold (i.e., shocked) misses were analyzed.

For each perceptual training session, percent correct responses were plotted as a function of AM depth and fit with a cumulative Gaussian function using the open-source toolbox psignifit 4^104^. We clamped the fit to the lowest AM depth presented (options.stimulusRange = [lowest AM, -0.01]); otherwise, default parameters were used. After fitting, data were converted back to *d’* space. Thresholds were defined as the AM depth at which the fitted curve crossed *d’* = 1.

To identify learners and non-learners, we quantified performance using two metrics. First, improvement was calculated as the difference between the threshold on the first day of perceptual training and the average threshold across the final three training days. Second, each subject’s threshold was plotted as a function of log_10_(day), and the slope of a linear regression fit to these data was used as a measure of the rate and direction of threshold change. Animals exhibiting either negative improvement or a positive regression slope, both indicative of worsening perceptual performance, were classified as non-learners and excluded from analyses of learning-related changes (Figure S1A).

Neural measures (described below) were compared between early (day 1) and late (days 8-10) stages of perceptual training. Late training was defined as the period after perceptual performance had reached asymptote. To estimate the timing of asymptote, mean AM detection thresholds were plotted as a function of training day and fit with an exponential decay^105–108^. The fitted curve approached a final asymptotic threshold of -16.51 dB and reached 95% of the total threshold improvement by training day 7.6 (Figure 1E). Accordingly, days 8–10 were classified as late-stage (asymptotic) training.

### Electrode implant procedures

NeuroNexus 64-channel probes (A4x16-Poly2-5mm-20s-lin-160) were mounted on in-house 3D-printed microdrives. Just before implantation, the backs of the electrode shanks were painted with fluorescent microspheres (FluoSpheres 0.2 µm; Invitrogen) that were dialyzed for sodium azide removal using a dialysis cassette (Slide-A-Lyzer 10K; ThermoFisher) soaked overnight in PBS at 4 °C.

One day before surgery, minocycline (0.02 mg/mL) was added to the drinking water and meloxicam (1.5 mg/kg) was administered subcutaneously. On the day of the surgery, animals were anesthetized with isoflurane (5%) in medical grade oxygen (>93%) in an induction chamber. The head was shaved, and a second dose of meloxicam plus dexamethasone (0.35 mg/kg) was delivered subcutaneously. The animal was placed on a warming pad, and the head was secured in a stereotaxic frame (Kopf). Isoflurane (1-3%) and oxygen (>93%) were steadily delivered at a flow rate of 2 L/min throughout the surgery. Lubricant ophthalmic ointment was applied to the eyes to prevent desiccation. The top of the head was sanitized with alternating swabs of 70% isopropyl alcohol and betadine. The skull was exposed, dried with hydrogen peroxide, and leveled between lambda and bregma. Four bone screws were inserted, craniotomy locations were marked, and the skull was covered with C&B Metabond (Parkell) before craniotomies were performed. Electrodes were implanted using the following coordinates (all relative to lambda): 1.20 (medial shank) through 1.60 (lateral shank) mm lateral, 9.05-9.50 mm rostral, 2.50-3.20 mm ventral, angle=0° from vertical. Following electrode implantation, the craniotomy and electrode shanks were covered with KwikCast (WPI), all moving pieces—exposed electrode shanks, circuit board, flex cable, microdrive—were coated in petroleum jelly to prevent cementing, and the implant was secured and protected with dental cement (Palacos). Animals recovered on heat under observation and were provided with minocycline-treated drinking water for the following seven days. Controlled water access and behavioral experiments began at least seven days after surgery.

### Electrophysiology acquisition and processing

Electrophysiological activity was recorded extracellularly from the OFC of freely-moving animals during behavioral training sessions that took place every 24-48 hours. Signals were collected using one of two acquisition systems: (I) Analog signals were amplified and digitized at 24.4 kHz (PZ5; Tucker Davis Technologies, TDT) via a 64-channel wireless headstage and receiver (W64, Triangle BioSystems), and synchronized with behavioral timestamps through RZ2 and RZ6 processors running the Synapse software suite (TDT) and the Synapse API for MATLAB (MathWorks). Raw signals were saved using an RS4 data streamer (TDT). (II) Signals were amplified and digitized at 30 kHz by a 64-channel RHD headstage (Intan Technologies) connected to a slip-ring commutator (Taidacent; purchased from Amazon), and synchronized with behavioral timestamps via an RHD recording controller (Intan) and OpenEphys software^109^.

Offline signals were high-pass filtered at 150 Hz and common-median referenced across channels. Spike sorting was performed using Kilosort4^110^. Sorting results were manually curated using Phy (https://github.com/cortex-lab/phy). During curation, clusters were classified as putative single units based on a high signal-to-noise ratio, a clear refractory period gap in the autocorrelogram, and cross-correlogram isolation from neighboring clusters. Following curation, unit quality was assessed using Allen Brain Institute metrics (https://allensdk.readthedocs.io/en/latest/_static/examples/nb/ecephys_quality_metrics.html): ’presence ratio’, ’amplitude cutoff fraction missing’, and ’ISI violation false-positive rate’ (isi_threshold = 1.5 ms; min_ISI = 0.15 ms^111^). Additionally, a ’refractory period violation rate’ was computed as the fraction of interspike intervals falling within a 1.5 msec refractory period. Units were retained as true single units if they met the following criteria: presence ratio > 0.7, amplitude cutoff fraction missing < 0.2, ISI violation false-positive rate < 0.5, and refractory period violation rate < 2%.

Following unit isolation, firing rates were computed across three temporal windows: (1) Non-AM: a 0.9-sec window during correct rejection trials, when the animal was drinking from the spout prior to AM onset; (2) AM: a 0.9-sec window following AM stimulus onset; (3) Outcome: a 0.9-sec window following spout offset. Because miss trials were accompanied by a shock artifact spanning approximately ±0.3 sec around spout offset, the response window was shifted by 0.3 sec on miss trials to exclude the artifact.

To classify units by spike-waveform shape, we analyzed mean extracellular waveforms from well-isolated single units at the recording channel with largest amplitude. From each waveform we extracted three features (Figure S2B). Peak-to-peak duration was the time between the principal peak and the subsequent smaller peak. The peak-to-peak ratio was the absolute ratio of the first and second peak amplitudes. Repolarization duration^112^ was the time from the second peak to the subsequent inflection point, defined as the first zero-crossing of the second derivative of the waveform occurring after the second peak. Units with atypical waveforms—those whose principal trough amplitude was smaller than the subsequent peak—were excluded. These three features were z-scored and submitted to k-means clustering (k = 2, 25 random initializations). Only regular-spiking units (cluster with higher peak-to-peak duration; putative excitatory neurons) were analyzed in this study.

Tonic spike counts (i.e., during the non-AM period while animals quietly drank from the spout) showed substantial overdispersion relative to a Poisson process (Goodness of Poisson fit Akaike Information Criterion (AIC)= 4 x 10^3^) and were better described by a negative binomial distribution (AIC = 2.9 x 10^3^; Likelihood ratio test of Poisson versus binomial fits: χ²(1) = 1.1 x 10^3^, p < 0.001), characteristic of cortical neurons^113–115^. Therefore, firing rates were natural log-transformed [ln(FR + 1); the +1 offset avoids ln(0)] to stabilize variance across the population prior to baseline subtraction^116^, yielding a log-ratio of evoked to baseline (non-AM) firing. This baseline-normalized log ratio is henceforth referred to as ’normalized firing’.

Extracellular recording data from three animals in this study were included in a previously published manuscript^62^ that examined the contribution of OFC to task-dependent plasticity in the auditory cortex. That study did not include any analyses of perceptual learning, and all learning-related analyses presented here are therefore new. For the present manuscript, all data were reprocessed from scratch using Kilosort4 (rather than Kilosort2), and all analyses are novel.

### Neurometric threshold computation

To enable assessments of AM depth sensitivity^19,97,117^, neuronal firing rates were transformed into the signal detection index (*d’*) computed as:

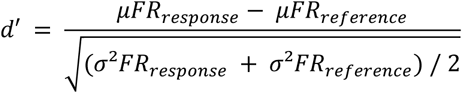

where μFR and σ^2^FR are the mean and variance of response and reference firing rate distributions, respectively.

To assess AM depth sensitivity on miss trials (Figure S2F-G), the response distribution consisted of log-transformed firing rates during the AM period, and the reference distribution consisted of log-transformed firing rates during the immediately preceding non-AM period.

To assess AM depth sensitivity during the spout offset period (Figure 2F-H), the response distribution consisted of normalized firing rates (baseline-subtracted, log-transformed; see above) during the post-spout offset period. The reference distribution consisted of normalized firing rates evoked by the lowest AM depth presented during that session that had a sufficient trial count (≥3). The lowest AM depth was used as a reference because it was expected to contain minimal AM-related information while still including spout offset-related activity. In contrast, the preceding non-AM period lacks spout offset-related responses, and therefore does not provide an appropriate reference for assessing AM depth-dependent modulation of spout offset activity.

For each unit, neural *d’* values were sorted by AM depth and fit with a two-parameter sigmoid function of the form:

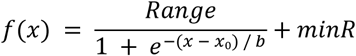

Where Range is the range of *d’* values, minR is the minimum *d’* value, and x is the AM depth. The parameters to be estimated were x₀ (the sigmoid inflection point) and b (the slope). The model was fit using nonlinear least squares (nlsLM), with the sign of b initialized from the direction of a Pearson correlation between AM depth and *d’*, and constrained so that x₀ remained within 20% of the depth range.

Previous studies defined neurometric threshold as the AM depth at which the neurometric fit crossed a *d’* of 1^19,97,117^. These studies focused on auditory cortical recordings and pooled hit and miss trials when calculating neural *d’*. In contrast, our analyses were restricted to either hit or miss trials, substantially reducing the number of trials contributing to each neural *d’* estimate. Because the firing rate variance inversely scales with the number of observations, and because we were recording in a higher-order brain region rather than a classical auditory structure, we expected to observe lower *d’* values than those reported previously. To account for this difference, we chose a more liberal criterion of *d’* = 0.5 when defining neurometric threshold. Units exhibiting monotonic fits that significantly differed from a flat line and reached an absolute *d’* value greater than 0.5 were therefore classified as AM depth sensitive.

### Permutation test for task-evoked firing rate modulation

To identify units responsive to trial outcome, we employed a permutation test comparing log-transformed firing rates during the outcome period to those during the preceding non-AM period on a trial-by-trial basis. For each unit, the observed test statistic was the difference in mean firing rates between the outcome and non-AM periods. A null distribution was generated by randomly permuting the period labels (outcome vs. non-AM) across all trials 999 times and recomputing the mean difference on each iteration, yielding a distribution of differences expected under the null hypothesis of no outcome-dependent modulation. A two-tailed p-value was computed as the proportion of permuted differences whose absolute value equaled or exceeded the absolute observed difference. This test was performed separately for each unit × trial type combination. Units with insufficient data (mean log firing rate < 0.1, fewer than 2 trials, or zero variance in paired differences) were excluded and classified as unresponsive.

### Fiber photometry

Pre-operative care and initial surgical steps were similar to electrode implant surgeries. AAV1-hSyn-axon-jGCaMP8s-P2A-mRuby3^118^ (GENIE Project; Addgene plasmid #172921) was injected into the left OFC to drive expression of a genetically-encoded calcium indicator (jGCaMP8s). A single 200 nL injection was made at the following coordinates relative to lambda: 1.2 mm lateral, 9.05 mm rostral, 3.20 mm ventral from pial surface, angle: 0° from vertical. Following the injection, the craniotomy was covered with KwikSil (WPI).

For auditory cortex (AC) fiber implants, the temporalis muscle insertion was detached and the muscle was separated from the skull using dry pieces of sterile SurgiFoam (Ethicon). A craniotomy was performed on the edge between the parietal and temporal bones. An optical fiber (400 µm core; Ø1.25 mm ceramic ferrule; 0.5 NA; RWD Life Science) was implanted in the left AC using the following coordinates relative to lambda: 5.7 mm lateral, 3.2 mm rostral, 1.7 mm ventral from pial surface; angle: 10° clockwise from vertical plane facing the animal’s posterior side. For visual cortex (VC) implants, procedures were similar to AC, except temporalis muscle detachment was not required. A craniotomy was performed over the parietal bone. An optical fiber was implanted in the left VC using the coordinates (relative to lambda): 2-4 mm lateral, 0 mm rostral, 0.55-0.7 mm ventral from pial surface; angle: 0° from vertical.

The craniotomy around the fiber was covered with KwikSil and the fiber was secured with dental cement. Post-operative care was as described above. Controlled water access and behavioral shaping started at least three weeks after surgery to allow for virus expression. Fiber photometry recordings began at least four weeks after surgery.

All optical components — including patch cords, fluorescence MiniCube, 405 and 465 nm connectorized LEDs, LED driver, and Newport photoreceiver module—were obtained from Doric Lenses. LED modulation and photovoltaic signal acquisition were performed at 1 kHz using an RZ2 processor and Synapse software (Tucker-Davis Technologies).

Calcium-dependent fluorescence was recorded with a 465 nm LED. A 405 nm LED was used simultaneously to capture calcium-independent isosbestic signals for correction of movement artifacts and photobleaching. The isosbestic point of GCaMP is ∼410 nm^119^.

Prior to each recording session, patch cords were photobleached overnight. Fiber-optic cannulas were coupled to the system via a quick-release connector (ADAL3; ThorLabs), and light intensity at the tip of a bare implantable fiber was measured using a photodiode power sensor and energy meter (S120C and PM100USB; ThorLabs). Light output was ∼200-325 and ∼25 µW (465 and 405 nm, respectively).

Photometry signals were processed offline similarly to^120,121^ using a custom MATLAB pipeline. Raw 465 and 405 nm signals were low-pass filtered at 3 Hz using a zero-phase Butterworth filter^120^. Recording segments contaminated by LED onset artifacts or movement noise were excluded based on time ranges selected manually. Baseline drift was corrected independently for each signal using the adaptive iteratively reweighted Penalized Least Squares algorithm (airPLS^122^; parameters: λ = 10⁷, iterationNumber = 1, threshold = 0.1, d = 0.5, maxIteration = 50). Both signals were then standardized by median subtraction and division by the standard deviation. The standardized 405 nm signal was fit to the standardized 465 nm signal using iteratively reweighted least squares regression (IRLS^120^; tuning constant = 1.4), and the resulting fit was subtracted from the 465 nm signal to yield the dF/F trace. This dF/F signal was then aligned to behavioral event timestamps and z-scored relative to a 0.25 sec pre-stimulus baseline.

### Histology and imaging

Upon completion of all experiments, each animal was deeply anesthetized with ketamine (150 mg/kg) and xylazine (6 mg/kg) in sterile saline and perfused transcardially, first with ∼20 mL phosphate-buffered saline (PBS) and then with ∼20 mL 4% paraformaldehyde (PFA) in PBS. Implants were subsequently removed, and each brain was harvested and immersed in 4% PFA for 1–7 days of post-fixation before sectioning. Brains were then returned to PBS, embedded in 6% agar, and cut at 70 µm on a vibratome (Leica VT-1000S). The resulting sections were mounted onto gelatin-coated slides and coverslipped with ProLong Gold or Diamond mountant (Molecular Probes).

Tissue expressing GFP (photometry experiments) was processed for GFP immunofluorescence to amplify the signal. Free-floating sections were first rinsed in PBS (3 × 10 min), then blocked for 2 h at room temperature in 10% normal goat serum (NGS) made up in PBS containing 0.3% Triton X-100 (Sigma; PBT). Sections were next transferred to rabbit anti-GFP primary antibody (1:1000; Invitrogen #A11122) diluted in the same NGS/0.3% PBT blocking solution, where they were held for 1 h at room temperature followed by 40–48 h at 4 °C. After three 15-min washes in 0.1% PBT, sections were incubated for 1 h at room temperature in goat anti-rabbit Alexa-488 secondary antibody (1:1000; Invitrogen #A32731). Finally, sections were given three final 10-min washes in 0.1% PBT and coverslipped with ProLong Diamond.

Imaging was performed at 10× and 40× on either an epifluorescence microscope (Leica DM750) or a confocal microscope (Zeiss LSM 980 with Airyscan 2). For epifluorescence acquisition, tiles were captured manually and merged into a single composite using the PhotoMerge panorama function in Photoshop 2023 (Adobe). All following image processing was done in ImageJ^123^.

## QUANTIFICATION AND STATISTICAL ANALYSIS

All data were processed offline by custom MatLab pipelines (https://github.com/caraslab, repositories: caraslab-behavior-analysis, caraslab-spikesortingKS4, and caraslab-fiberphotometry), then further analyzed by custom Python code (https://github.com/caraslab, repositories: Caraslab_FP_preprocessing_pipeline and Caraslab_EPhys_preprocessing_pipeline).

Statistical analyses were performed using R (version 4.5.3) and RStudio (version 2026.05.0+218; Posit Software). All statistical results can be found in Tables 1 and S1. Analyses were performed by generalized linear models followed by ANOVA (GLM/ANOVA) using the ‘glmmTMB’^124^ and the ‘car’^125^ packages for R. Normality of GLM residuals was assessed after each fit by visually inspecting the distribution of studentized residuals (q-q plots) using the ‘DHARMa’ package for R^126^. When normality was not fulfilled, data were ‘rankit’-transformed^127^, and GLMs were rerun. When interactions were significant, Bonferroni-adjusted Wald z-tests for multiple comparisons using the package ‘emmeans’ for R (https://github.com/rvlenth/emmeans) were used.

Preliminary analyses revealed no effect of sex on our main measures of interest, so data from males and females were always analyzed together.

## KEY RESOURCES TABLE

| REAGENT or RESOURCE | SOURCE | IDENTIFIER |
| --- | --- | --- |
| **Antibodies** |  |  |
| Rabbit polyclonal anti-GFP | Invitrogen | #A11122 |
| Goat anti-rabbit Alexa-488 | Invitrogen | #A32731 |
| **Bacterial and virus strains** |  |  |
| AAV1-hSyn-axon-jGCaMP8s-P2A-mRuby3 | Addgene | 172921-AAV1 |
| **Chemicals, peptides, and recombinant proteins** |  |  |
| FluoSpheres (0.2 μm) | Invitrogen | F-8848 |
| **Experimental models: Organisms/strains** |  |  |
| Gerbil (*Meriones unguiculatus*) | Charles River Laboratories | www.criver.com |
| **Software and algorithms** |  |  |
| Behavioral control software | ePsych (Dr. Daniel Stolzberg) | www.github.com/caraslab/epsych |
| Electrophysiology processing | Custom code; Kilosort 4<sup>110</sup>; Phy | www.github.com/caraslab/caraslab-spikesortingKS4<br>www.github.com/cortex-lab/phy |
| Photometry processing | Custom code | www.github.com/caraslab/caraslab-fiberphotometry |
| Behavioral data processing | Custom code | www.github.com/caraslab/caraslab-behavior-analysis |
| ImageJ | Schneider et al.<sup>123</sup> | www.imagej.nih.gov/ij |
| PyCharm | JetBrains | www.jetbrains.com |
| RStudio | Posit | www.posit.co |
| MatLab | MathWorks | www.mathworks.com |
| TDT Synapse | Tucker Davis Technologies | www.tdt.com |
| Open Ephys GUI | Open Ephys | www.open-ephys.org |
| Adobe Photoshop and Illustrator 2026 | Adobe | www.adobe.com |

## Notes

### Competing Interest Statement

The authors have declared no competing interest.

