## Supplementary text for "Perceptual training selectively strengthens top-down signaling to sensory cortex"

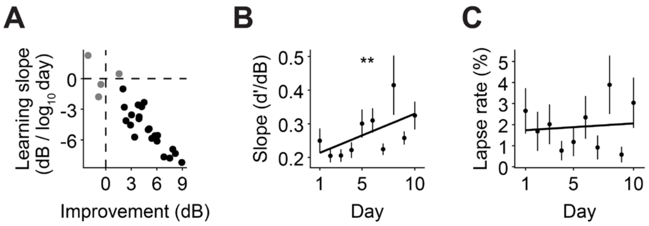


**Figure S1. Perceptual learning criterion, and slope and lapse rates. Related to Figure 1 and Table S1 (A)** Learners and non-learners were distinguished using two metrics: improvement (first-day threshold minus the mean threshold of the final three training days) and the slope of a linear fit to threshold versus $\log_{10}$(day). Subjects with negative improvement or a positive slope—both indicating worsening performance—were classified as non-learners and excluded from learning-related analyses. In total, four subjects (three females) were removed from learning-related analyses (grey points). **(B)** Behavioral slopes significantly improve across training. **(C)** Lapse rates remain low and unchanged across learning. $^{**}p < 0.01$.

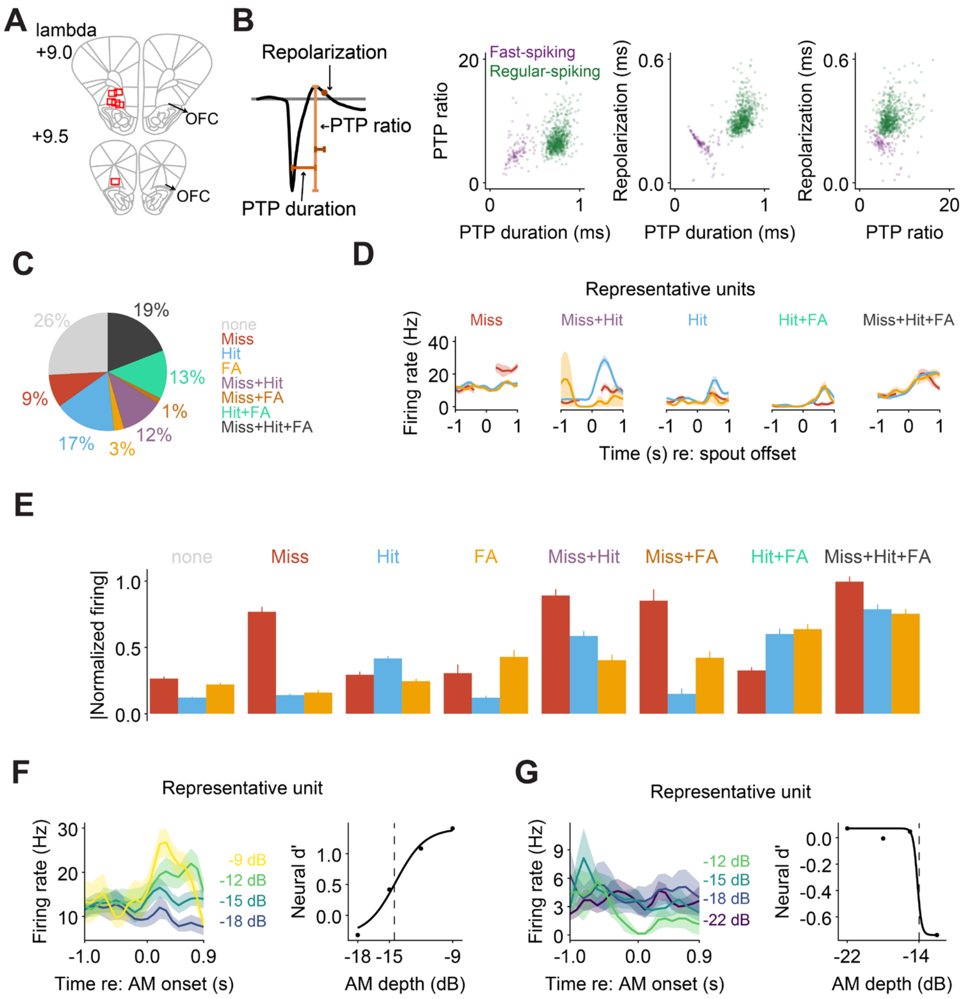


**Figure S2. OFC electrode placements, waveform classification, detailed trial outcome responsiveness, and sound-evoked responses in OFC single-units. Related to Figure 2 and Table S1** **(A)** Electrode placements were confirmed histologically post-hoc via fluorescently labeled tracks and/or electrolytic lesions (see Figure 2A). Red boxes represent final electrode locations from six animals. Electrodes were advanced ≤ 280 µm during all experiments. **(B)** Neuronal waveform classification parameters. Left: Three measurements used for *k*-means clustering (*k* = 2): peak-to-peak (PTP) ratio, PTP duration, and repolarization time[1]. Right: Pairwise correlation plots between the three measurements across OFC single-units. Colors illustrate classification results into fast- (purple) or regular-spiking (green) units. Only regular-spiking neurons were analyzed for this study. **(C)** Pie chart representing the proportion of units responding to trial outcome combinations as assessed via permutation tests measuring firing rates 0-1 sec after spout offset responses. Units unresponsive to any trial types were classified as ‘none’. FA: false alarm. **(D)** Raw firing rates for five representative units that responded to distinct combinations of trial outcome types. **(E)** Mean ± SEM absolute normalized firing from units responsive to each outcome combination group. **(F)** Left: raw firing rate of a representative unit to a range of AM depths, aligned to AM onset. Only miss trial activity is shown. This unit exhibited increased firing as the AM depth increased. Right: neural *d’* values (referenced to the preceding non-AM noise) plotted as a function of AM depth and fit with a sigmoid. The dashed line illustrates this unit’s threshold (where the sigmoid crosses *d’* = 0.5; see STAR methods). **(G)** Same as in (F) but for a unit that exhibited decreased firing as AM depth increased.

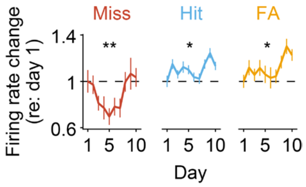


**Figure S3. Perceptual training selectively strengthens trial outcome responses in OFC neurons. Related to Figure 3 and Table S1.** For each trial outcome and subject, the normalized post spout-offset period firing on day 1 was averaged across all units within a subject and set as reference. Firing rate change was computed as log-ratios relative to that baseline. Across training, firing rates on hit and false alarm trials increased, while firing rates on miss trials exhibited a transient reduction followed by renormalization. *p < 0.05; **p < 0.01.

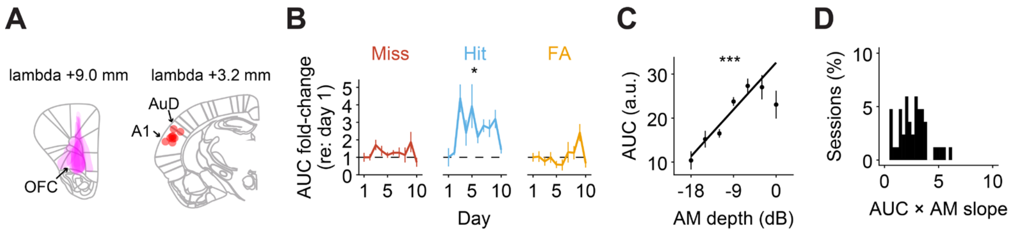


**Figure S4. Perceptual training selectively strengthens outcome signaling in the OFC-to-AC pathway. Related to Figure 5 and Table S1. (A)** Left: AAV1-axon-jGCaMP8s injection location and spread in OFC was estimated for each animal from the distribution of mRuby3-expressing neurons. Right: Fiber tip locations in AC for each subject. A1 = primary auditory cortex, AuD = dorsal auditory cortex. **(B)** Training day selectively predicts stronger OFC-to-AC axonal calcium responses on hit trials. For each trial outcome and subject, the area under the curve (AUC; 0-3 sec) on day 1 was averaged and set as reference. Normalized AUC change was computed as ratios to that baseline. During training, AUC on hit trials increased, while AUC values during miss and false alarm trials did not change. **(C)** Mean ± SEM AUC across all sessions is strongly correlated with AM depth. **(D)** Distribution of slopes from sessions with significant Spearman correlations between AUC and AM depth (51% of all sessions). *$p < 0.05$. ***$p < 0.001$

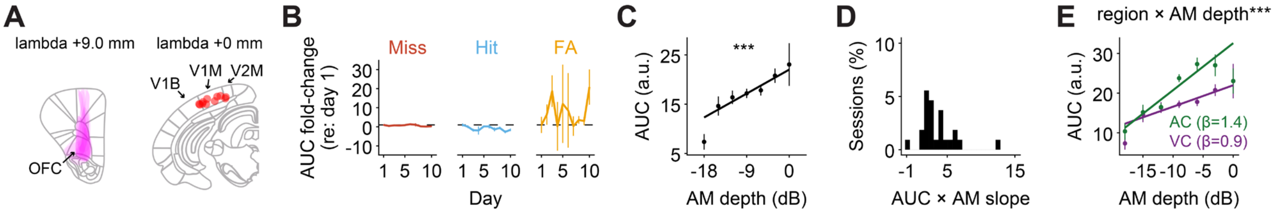


**Figure S5. Auditory perceptual training does not alter signaling in the OFC-to-visual cortex pathway. Related to Figure 6, Figure S4C and Table S1. (A)** Left: AAV1-axon-jGCaMP8s injection location and spread in OFC was estimated for each animal from the distribution of mRuby3-expressing neurons. Right: Fiber tip locations in VC for each subject. V1B = primary visual cortex, binocular area, V1M = primary visual cortex, monocular area, V2M = secondary visual cortex, medial area. **(B)** Training day has no effect on OFC-to-VC calcium responses for any trial outcome. For each trial outcome and subject, the area under the curve (AUC; 0-3 sec) on day 1 was averaged and set as reference. Normalized AUC change was computed as ratios to that baseline. **(C)** Mean ± SEM AUC across all sessions is strongly correlated with AM depth. **(D)** Distribution of slopes from sessions with significant Spearman correlations between AUC and AM depth (23% of all sessions). **(E)** Comparison of AUC and AM depth relationship in the OFC-to-AC and OFC-to-VC pathway. Data are mean ± SEM AUC and AM depth across all sessions. Correlation slopes (β; indicated on the plot) were significantly higher in AC than in VC. OFC-to-AC data are replotted from Figure S4C. ***$p < 0.001$.

**Table S1.** Supplementary statistical results

| Figure | Test | Predictor(s) | Statistic | P value |
|---|---|---|---|---|
| S1B | GLM/ANOVA | Day | $\chi^2_1 = 8.074$ | **0.004** |
| S1C | GLM/ANOVA | Day | $\chi^2_1 = 0.123$ | 0.725 |
| S3 | GLM/ANOVA: Miss | poly(Day, 2) | $\chi^2_1 = 11.099$ | **0.004** |
| | | AM depth (covariate) | $\chi^2_1 = 0.280$ | 0.597 |
| | GLM/ANOVA: Hit | poly(Day, 2) | $\chi^2_1 = 8.382$ | **0.015** |
| | | AM depth (covariate) | $\chi^2_1 = 0.011$ | 0.916 |
| | GLM/ANOVA: FA | poly(Day, 2) | $\chi^2_1 = 8.470$ | **0.015** |
| S4B | GLM/ANOVA: Miss | Day | $\chi^2_1 = 2.514$ | 0.113 |
| | | AM depth (covariate) | $\chi^2_1 = 0.633$ | 0.426 |
| | GLM/ANOVA: Hit | Day | $\chi^2_1 = 5.127$ | **0.024** |
| | | AM depth (covariate) | $\chi^2_1 = 115.122$ | **< 0.001** |
| | GLM/ANOVA: FA | Day | $\chi^2_1 = 0.005$ | 0.944 |
| S4C | GLM/ANOVA | AM depth | $\chi^2_1 = 229.115$ | **< 0.001** |
| S5B | GLM/ANOVA: Miss | Day | $\chi^2_1 = 1.383$ | 0.239 |
| | | AM depth (covariate) | $\chi^2_1 = 1.004$ | 0.316 |
| | GLM/ANOVA: Hit | Day | $\chi^2_1 = 2.909$ | 0.088 |
| | | AM depth (covariate) | $\chi^2_1 = 2.100$ | 0.147 |
| | GLM/ANOVA: FA | Day | $\chi^2_1 = 0.264$ | 0.607 |
| S5C | GLM/ANOVA | AM depth | $\chi^2_1 = 68.305$ | **< 0.001** |
| S5E | GLM/ANOVA | Region | $\chi^2_1 = 8.266$ | **0.004** |
| | | AM depth | $\chi^2_1 = 287.472$ | **< 0.001** |
| | | Interaction | $\chi^2_1 = 33.240$ | **< 0.001** |

GLM: Generalized linear model; ANOVA: Analysis of Variance; FA: False alarm; Bolded p-values indicate statistical significance.
